# piClusterBusteR: Software for Automated Classification and Characterization of piRNA Cluster Loci

**DOI:** 10.1101/133009

**Authors:** Patrick Schreiner, Peter W. Atkinson

**Affiliations:** Interdepartmental Graduate Program in Genetics, Genomics & Bioinformatics, University of California, Riverside, CA 92521, USA; Department of Entomology and Institute for Integrative Genome Biology, University of California, Riverside, CA 92521, USA

## Abstract

**Background:** Piwi-interacting RNAs (piRNAs) are sRNAs that have a distinct biogenesis and molecular function from siRNAs and miRNAs. The piRNA pathway is well-conserved and shown to play an important role in the regulatory capacity of germline cells in Metazoans. Significant subsets of piRNAs are generated from discrete genomic loci referred to as piRNA clusters. Given that the contents of piRNA clusters dictate the target specificity of primary piRNAs, and therefore the generation of secondary piRNAs, they are of great significance when considering transcriptional and post-transcriptional regulation on a genomic scale. A quantitative comparison of top piRNA cluster composition can provide further insight into piRNA cluster biogenesis and function.

**Results:** We have developed software for general use, piClusterBusteR, which performs nested annotation of piRNA cluster contents to ensure high-quality characterization, provides a quantitative representation of piRNA cluster composition by feature, and makes available annotated and unannotated piRNA cluster sequences that can be utilized for downstream analysis. The data necessary to run piClusterBusteR and the skills necessary to execute this software on any species of interest are not overly burdensome for biological researchers.

piClusterBusteR has been utilized to compare the composition of top piRNA generating loci amongst 13 Metazoan species. Characterization and quantification of cluster composition allows for comparison within piRNA clusters of the same species and between piRNA clusters of different species.

**Conclusions:** We have developed a tool that accurately, automatically, and efficiently describes the contents of piRNA clusters in any biological system that utilizes the piRNA pathway. The results from piClusterBusteR have provided an in-depth description and comparison of the architecture of top piRNA clusters within and between 13 species, as well as a description of annotated and unannotated sequences from top piRNA cluster loci in these Metazoans.

piClusterBusteR is available for download on GitHub: https://github.com/pschreiner/piClusterBuster

## Introduction

*p*-element induced wimpy testis (PIWI) proteins and the utilization of the PIWI-interacting RNA (piRNA) pathway has been conserved in a diverse range of Metazoans, including sponges, roundworms, fruit flies, and humans (Grimson et al. 2008). The importance of the role of piRNAs in fertility was demonstrated in Metazoans by the observation of crosses after exposure of *Drosophila melanogaster* to the *P* transposable element (TE) (Kidwell 1983). When female flies containing the *P* element were mated with males that were lacking it, the progeny were fertile. However, when males containing the *P* element were mated with females lacking it, hybrid dysgenesis occurred, leading to sterility of the progeny (Kidwell et al. 1977). It was later discovered that exposure to the *P* element prompted maternal deposition of piRNAs to effectively silence the *P* element and allow for fertile progeny (Brennecke et al. 2008).Perturbations to the piRNA pathway have also demonstrated gametogenic defects in *M. musculus* and *D. rerio* (Kuramochi-Miyagawa et al. 2004; Houwing et al. 2007). Although, piRNAs are notably absent in plant and fungal species (Grimson et al. 2008).

piRNAs are a subset of sRNAs between 24-31 nucleotides in length, although the range of the piRNA size distribution varies across species (Aravin et al. 2007; Arensburger et al. 2011). piRNAs are generated via a primary or secondary mechanism of biogenesis (Aravin et al. 2007).

Primary piRNAs derive from discrete genomic loci that are referred to as piRNA clusters. These loci can vastly range in size from under one thousand nucleotides to over one hundred thousand nucleotides in length. Transcription of piRNA clusters can occur in several distinct manners depending on the nature of the piRNA clusters (Brennecke et al. 2007).

piRNA clusters are characterized as unidirectional, bidirectional, or dual-stranded based on the direction transcription at the locus (Brennecke et al. 2007; Malone et al. 2009). The transcripts generated from piRNA clusters serve as precursor molecules for piRNAs, undergoing dicer-independent slicing and modification at their 3’ end (Aravin et al. 2007; Saito et al. 2007). The processing of primary piRNAs has been shown to demonstrate a bias of U at position 1 of the piRNAs (Brennecke et al. 2007). When post-transcriptional processing of piRNAs is complete, the molecules are referred to as mature piRNAs.

Mature piRNAs then associate with an Argonaute family, PIWI protein to form a RNA-induced silencing complex (RISC). The RISC complex has the capability to facilitate both transcriptional and post-transcriptional regulation. RISC-mediated transcriptional regulation occurs via piRNA association and guiding of a PIWI protein which facilitates epigenetic modification in *Drosophila* (Yin & Lin 2007). RISC-mediated post-transcriptional regulation occurs via piRNA association with PIWI or AGO3, which leads to piRNA-directed cleavage of mRNAs in *Drosophila* (Aravin et al. 2007). piRNAs have also been implicated in post-transcriptional silencing of mRNAs via poly(A) deadenylation (Rouget et al. 2010; Barckmann et al. 2015).

The number of PIWI proteins can differ in Metazoan species. While three PIWI proteins have been identified in *D. melanogaster, H. sapiens*, and *M. musculus*, as few as two PIWI proteins have been identified in *D. rerio* and as many as seven PIWI proteins have been identified in *Ae. aegypti* (Aravin et al. 2007; Keam et al. 2014; Kuramochi-Miyagawa et al. 2001; Houwing et al. 2007; Vodovar et al. 2012). It has not yet been determined whether the variation in the number of PIWI proteins between these species is a result of redundant, compensatory, or additional functionality.

Secondary piRNAs are generated by the slicing mechanism of RISC regulation, resulting in what is referred to as the amplification loop, or ping-pong pathway (Brennecke et al. 2007). The amplification loop functions by primary piRNA targeting of mRNA via sequence complementarity, followed by PIWI-mediated slicing of the target mRNA. The remaining fragment of the mRNA can be processed into a secondary piRNA. A secondary, mature piRNA can then associate with AGO3, and slice other mRNA targets via sequence complementarity in *Drosophila*. The overlap of complementarity between the piRNAs and their mRNA targets is generally ten base pairs in the opposite orientation, leaving an A10 bias in secondary piRNAs (Brennecke et al. 2007). piRNAs have been known to target TEs, genic mRNAs, viral mRNAs, and even rRNA molecules (Brennecke et al. 2007; Aravin et al. 2007; Yin & Lin 2007; Aravin et al. 2008; Garc’\ia-López et al. 2014).

Given that the contents of piRNA clusters dictate target specificity, finding the origin of these sequences is of great importance in understanding the biogenesis and function of piRNA clusters. We have developed software, piClusterBusteR, to be capable of automatically, consistently, and efficiently detecting top piRNA cluster loci and thoroughly describing the contents of those loci on a large scale. The capability that piClusterBusteR has to quantify piRNA cluster composition and describe annotated, as well as unannotated sequences in diverse Metazoan species allows for meaningful comparisons that can aid in facilitating a better understanding of top piRNA cluster biogenesis and function across species. Exploring piRNA cluster composition on a large scale can provide insight into conserved piRNA cluster architecture that dictates piRNA cluster biogenesis and function.

## Materials and Methods

piClusterBusteR is a series of integrated R and bash scripts that interact along with other standalone bioinformatics programs to perform piRNA cluster characterization and annotation. The tool supports a variety of user input data, customization of the analyses and computational resources to be used in executing the program. piClusterBusteR is intended to be executed in a Unix environment and has a series of required software dependencies (Table 2)

piClusterBusteR only requires four input parameters from the user on the command line: (1) input data, (2) a reference genome, (3) a species-specific gene set, and (4) a set of known TEs. Additional options are available to increase the efficiency of the software and to customize the program output.

piClusterBusteR allows for data input in the form of sRNA reads, piRNA cluster sequences, or piRNA cluster chromosomal loci. When sRNA reads are provided as the data input, piClusterBusteR must perform additional steps in order to assign piRNA cluster loci. First, all of the reads are filtered in order to analyze only those that are 24 nucleotides in length or greater. The piRNAs from the filtered FASTQ file are then mapped to the user-provided reference genome using proTRAC’s sRNA mapping tool. The piClusterBusteR-generated map file is then utilized to define the top piRNA cluster loci using proTRAC (Rosenkranz & Zischler 2012).

proTRAC is an standalone tool designed for the definition of piRNA clusters (Rosenkranz & Zischler 2012). proTRAC considers features of small RNA sequence reads features that are indicative of piRNAs such as read length and a U1 or A10 bias. proTRAC uses a density-based approach to identify genomic regions that have piRNA accumulation, as defined by a significant deviation from a hypothetical uniform distribution, which then defines the degree of confidence of the piRNA cluster call. proTRAC has demonstrated efficacy in piRNA cluster definition relative to previously established methods of piRNA cluster detection (Rosenkranz & Zischler 2012). The proTRAC output is then processed to identify the top piRNA cluster loci, as defined by the number of normalized reads per piRNA cluster, and converted to a BED file of piRNA cluster loci. The BED file is then utilized to analyze the contents of the individual top piRNA cluster loci.

In the individual piRNA cluster level analysis, piClusterBusteR performs a detailed characterization and quantification regarding the contents of each individual piRNA cluster. The user has the option to analyze piRNA clusters sequentially (default), or in parallel for each piRNA cluster of interest.

piClusterBusteR first extracts the sequence using the chromosomal coordinates and reference genome that was provided by the user. piClusterBusteR then attempts to identify the origin of the sequences within the piRNA clusters of interest.

In order to best infer the origin of the sequences within a given piRNA cluster, piClusterBusteR utilizes what we refer to as nested annotation using RepeatMasker, CENSOR, and BLAST (Smit et al. 1996; Jurka et al. 1996; Altschul et al. 1990). Nested annotation allows for sequential and non-redundant definition of known sequences with the piRNA cluster sequences under observation. RepeatMasker is run initially on the piRNA cluster of interest using the TE database and organism-specific gene set provided by the user (Smit et al. 1996). TE and organism-specific data sets were extracted from RepBase and NCBI non-redundant nucleotide databases, respectively (Jurka et al. 2005; Sayers et al. 2011). Any of the unannotated sequence remaining in the piRNA cluster of interest is extracted and subjected to TE and genic analysis via CENSOR (Jurka et al. 1996). Finally, the remainder of the unannotated piRNA cluster sequence is subjected to a blastn search, with a word size of 7 and maximum E-value of 1e-3, against the NCBI nucleotide database (Jurka et al. 1996; Altschul et al. 1990). Any of the hits returned in the BLAST of sequences within the NCBI non-redundant (nt) database are classified as “Other,” in comparison to sequence originating from known TEs or genes (Figure 3.2). Regions of the piRNA cluster loci that have not been defined with a known sequence origin are then extracted and printed reported. In doing so, piRNA cluster sequence of unknown origin can be easily accessed for downstream analysis.

To ensure that there is no redundancy in the sequence characterization, feature filtering is performed to only retain the best available annotation for a given piRNA cluster sequence. The best available annotation is defined as the hit with the longest available alignment length and highest similarity percentage to a known feature.

Non-redundant TE, genic, and “other” annotation is then summarized and plotted. A directory containing all of the intermediate annotation files and summary files is output in an individual directory to represent the analysis of each individual piRNA cluster. The results of the piRNA cluster level analyses of each piRNA cluster are stored so that they can be used in the genome level analysis.

In the genome level analysis, annotation is graphically compared between individual piRNA clusters. The piRNA clusters are compared in terms of their length, contents, degree of strand specificity, and percent genome occupancy. Top piRNA cluster loci can then be compared between piRNA clusters within the same species and between species on a genomic level (Figure 3.1).

**Figure 3.1.**
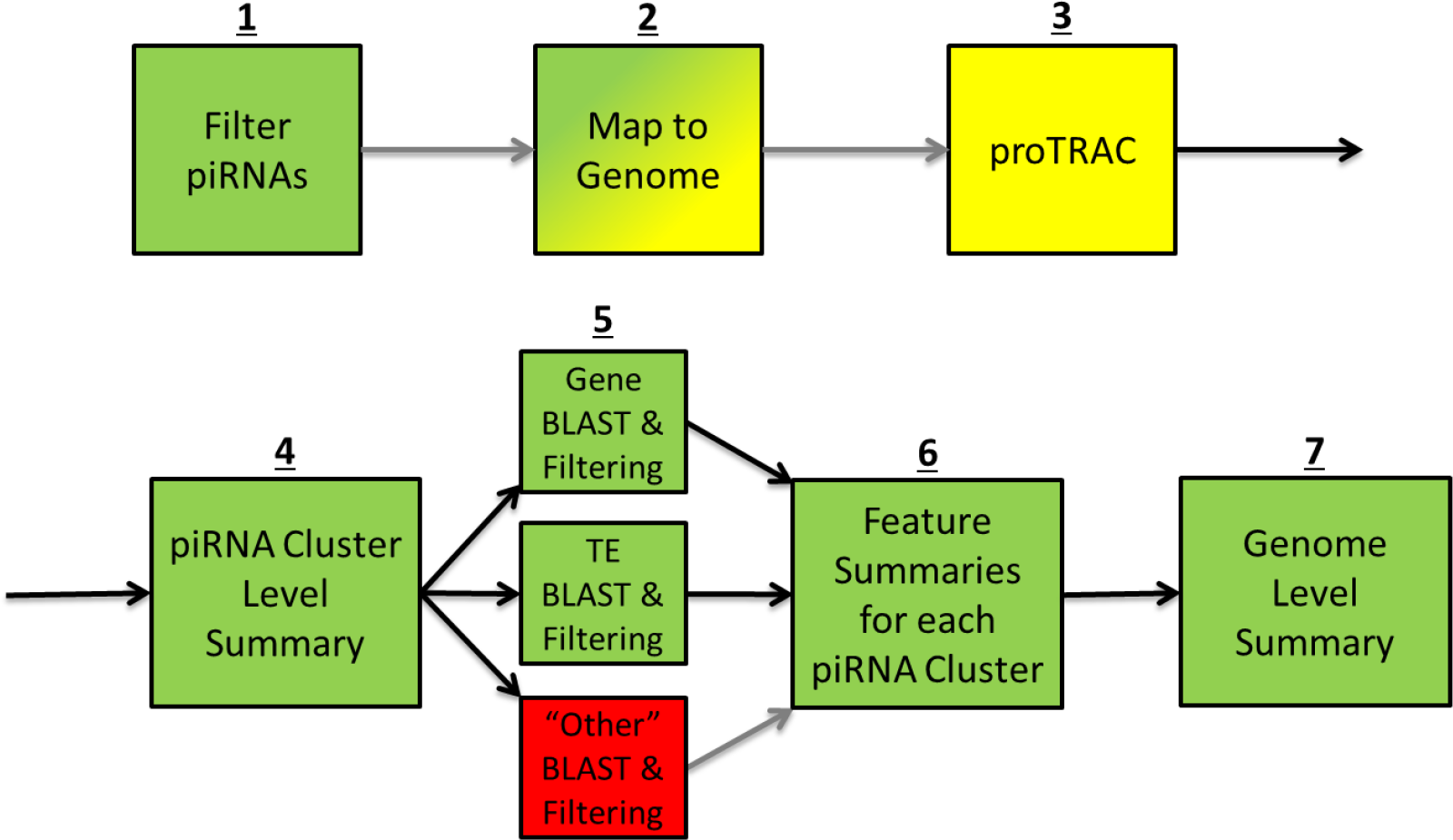
Algorithm Overview. A workflow of the steps taken by piClusterBusteR to annotate and characterize piRNA clusters. The step number is indicated above the boxes that describe the analysis in each step. The relative time requirement of each step is indicated by green, yellow, and red from fastest to slowest. The gray arrows indicate that the previous step may be skipped if it is unnecessary.

**Figure 3.2.**
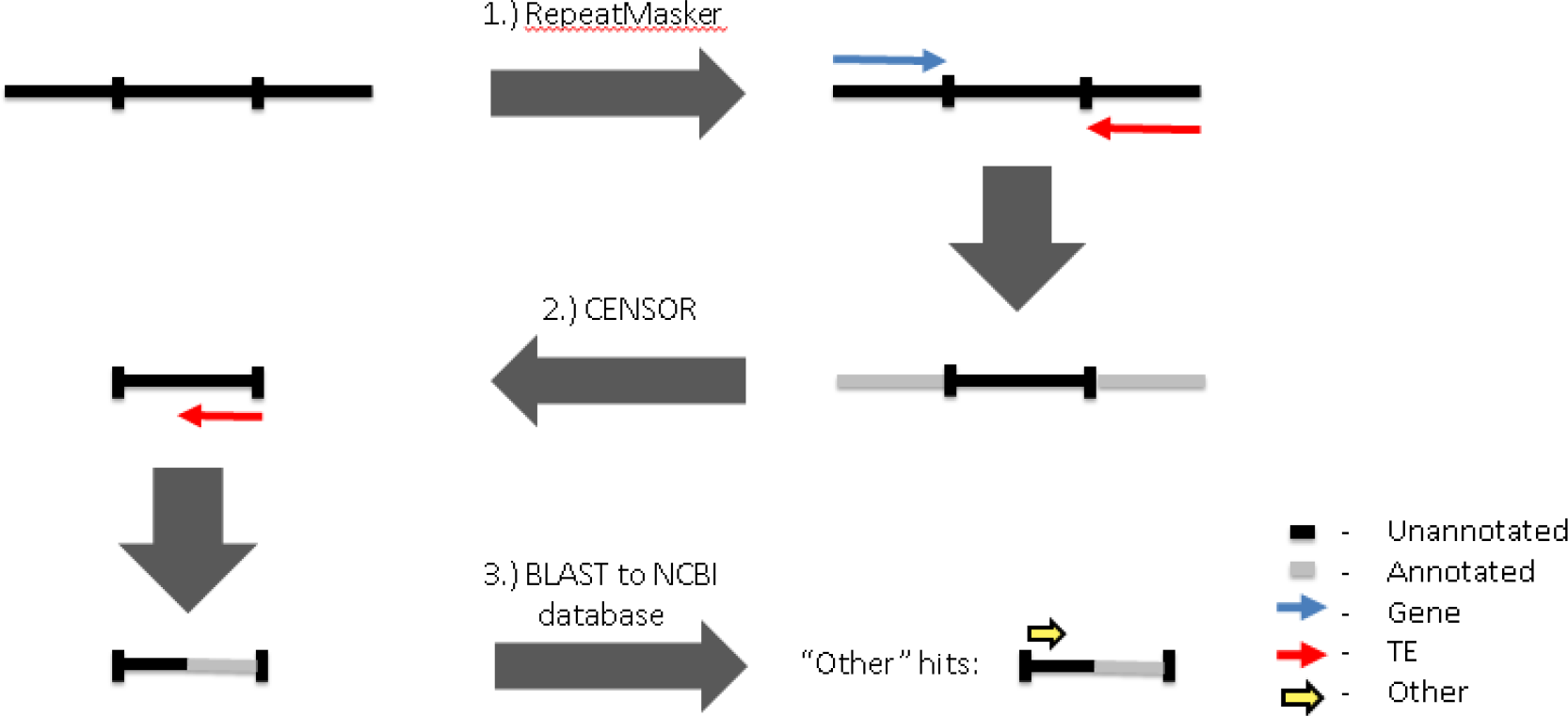
Nested Annotation. Workflow regarding the characterization of unannotated loci are using RepeatMasker, CENSOR, and BLAST (Smit et al. 1996; Jurka et al. 1996; Altschul et al. 1990). Sequence characterization in the former steps excludes sequences from being passed to the latter.

The main directory represents the outcome of the genome-level analysis. Four output files are generated in the genome-level analysis: (1) a BED file containing the piRNA cluster coordinates,(2) an aggregate file describing the total occupancy of piRNA clusters relative to the size of the organism’s genome, as well as the final data necessary to make the genome summary plots in a(3) graphical and (4) text format (Quinlan & Hall 2010). The genome-level graphical output contains a comparison of piRNA cluster size, piRNAs associated with each piRNA cluster, feature composition, and strandedness of feature calls, followed by the average feature content composition across all piRNA cluster loci analyzed (Figure 3.3).

**Figure 3.3.**
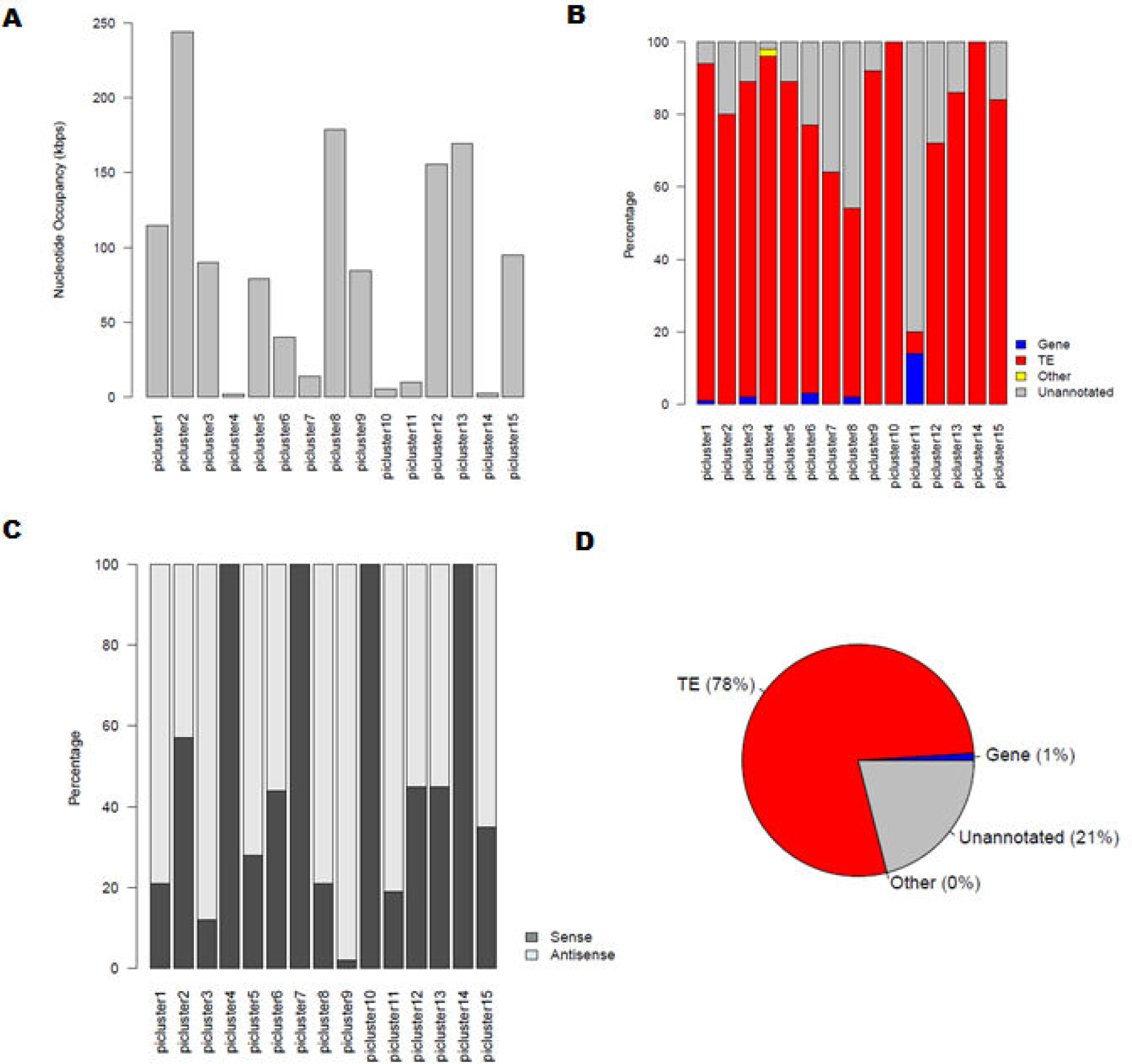
Genome-Level Analysis of Top 15 piRNA Cluster Contents in *Drosophila melanogaster*. **(A)** Comparison of piRNA clusters by the total number of nucleotides occupied **(B)** Relative nucleotide occupancy occupied by each feature **(C)** Stranded nucleotide occupancy of feature calls. Unannotated sequences are not considered in this representation **(D)** Average nucleotide occupancy occupied by each feature across the top 15 *D. melanogaster* piRNA clusters previously identified (Brennecke et al. 2007). The *flamenco* and *42AB* loci are represented as piRNA clusters 1 and 2, respectively (Aravin et al. 2007; Brennecke et al. 2007).

The genome level analysis also provides an individual directory for each piRNA cluster of interest in the order specified within the BED file of piRNA cluster loci. Within each piRNA cluster directory resides intermediate data files and summary files that are necessary to produce the piRNA cluster-level graphical output. The intermediate data files that were used in the data collection are available in the respective program output format defaults for each utilized tool (Table 1). The unfiltered BLAST output for each piRNA cluster, however, can often be large in size and is therefore removed by default. The piRNA cluster-level summary is also available in a text and graphical output.

**Table 3.1.**
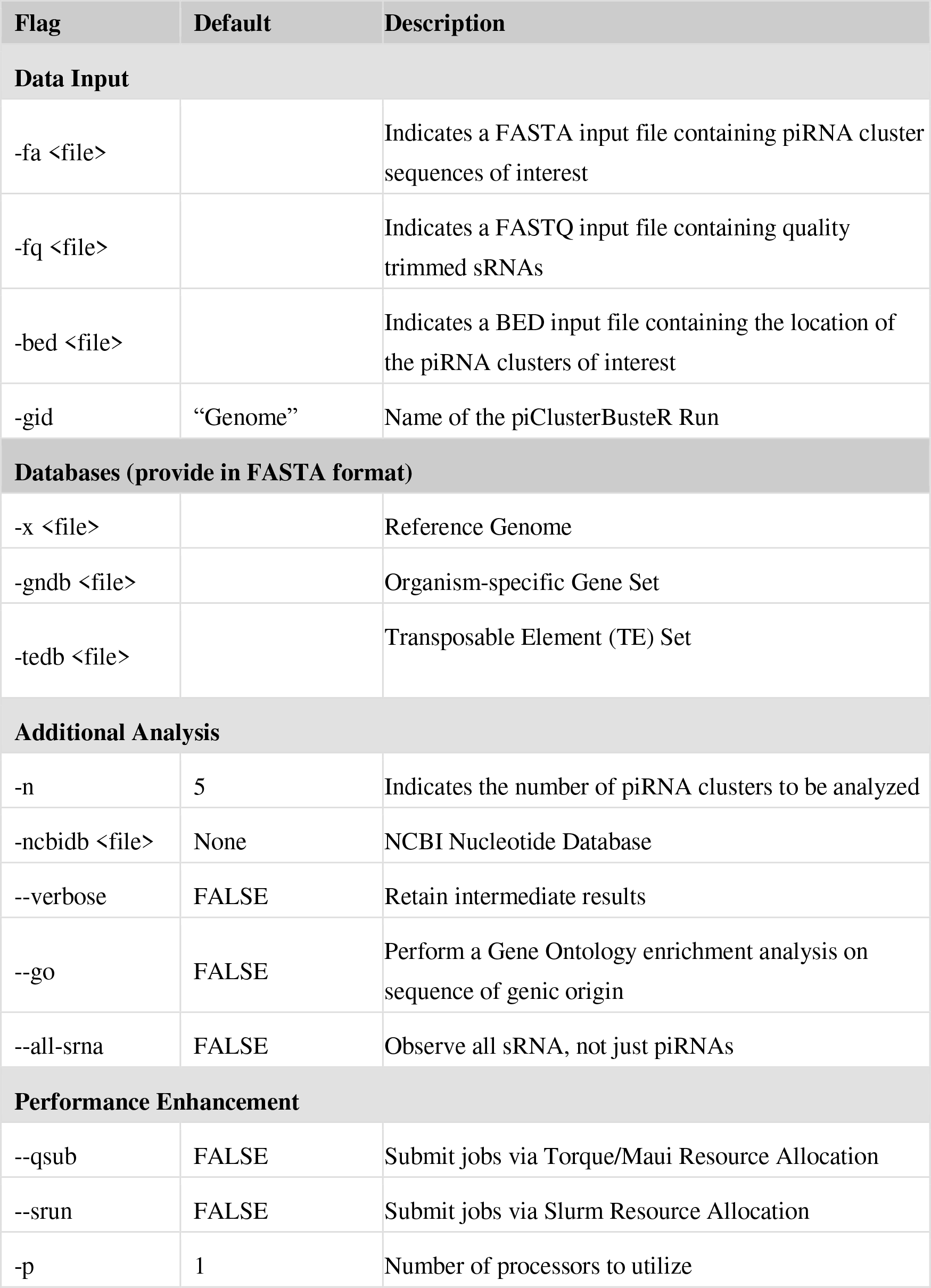
Program Parameters. A list of the options, corresponding runtime flags, and default values available for use in piClusterBusteR. A flag is an indicator to specify the type of input information to the application. A blank in the “Default” column constitutes a required parameter.

**Table 3.2.** List of Software and Databases Utilized in piClusterBusteR. Depending on the user-specified analyses to be performed, piClusterBusteR may require these **(A)**standalone software and **(B)** R libraries.

| Database | Website | Reference |
| --- | --- | --- |
| NCBI (nt) | http://www.ncbi.nlm.nih.gov/ | (Altschul et al. 1990) |
| RepBase | http://www.girinst.org/repbase/ | (Jurka et al. 2005) |

| Standalone Programs | Website | Reference |
| --- | --- | --- |
| BLAST+ (v2.2.30+) | http://blast.ncbi.nlm.nih.gov/ | (Sayers et al. 2011) |
| CENSOR | http://www.girinst.org/censor/ | (Jurka et al. 1996) |
| proTRAC | http://www.smallrnagroup.uni-mainz.de/ | (Rosenkranz & Zischler 2012) |
| RepeatMasker | http://www.repeatmasker.org/ | (Smit et al. 1996) |

| R Packages | Reference |
| --- | --- |
| Biostrings | (Pages et al. 2009) |
| doMC | (Analytics 2014) |
| GenomicRanges | (Lawrence et al. 2013) |
| gProfileR | (Reimand et al. 2007) |
| Plyr | (Wickham 2009) |
| Qcc | (Scrucca 2004) |
| Seqinr | (Charif & Lobry 2007) |
| systemPipeR | (Girke 2014) |

The piRNA cluster-level graphical output contains a representation of the number of each feature that was characterized within the piRNA cluster, the nucleotide occupancy of each feature called, the nucleotide occupancy of all feature calls in both orientations, a representation of the prominent TE superfamilies within the piRNA cluster, the prominent specific TEs called within the piRNA cluster, and optionally, the most significant GO terms associated with genic hits within the piRNA cluster, a GO enrichment analysis of genic hits within the piRNA cluster, and stranded sRNA coverage plot with annotated features (Figure 3.4).

**Figure 3.4.**
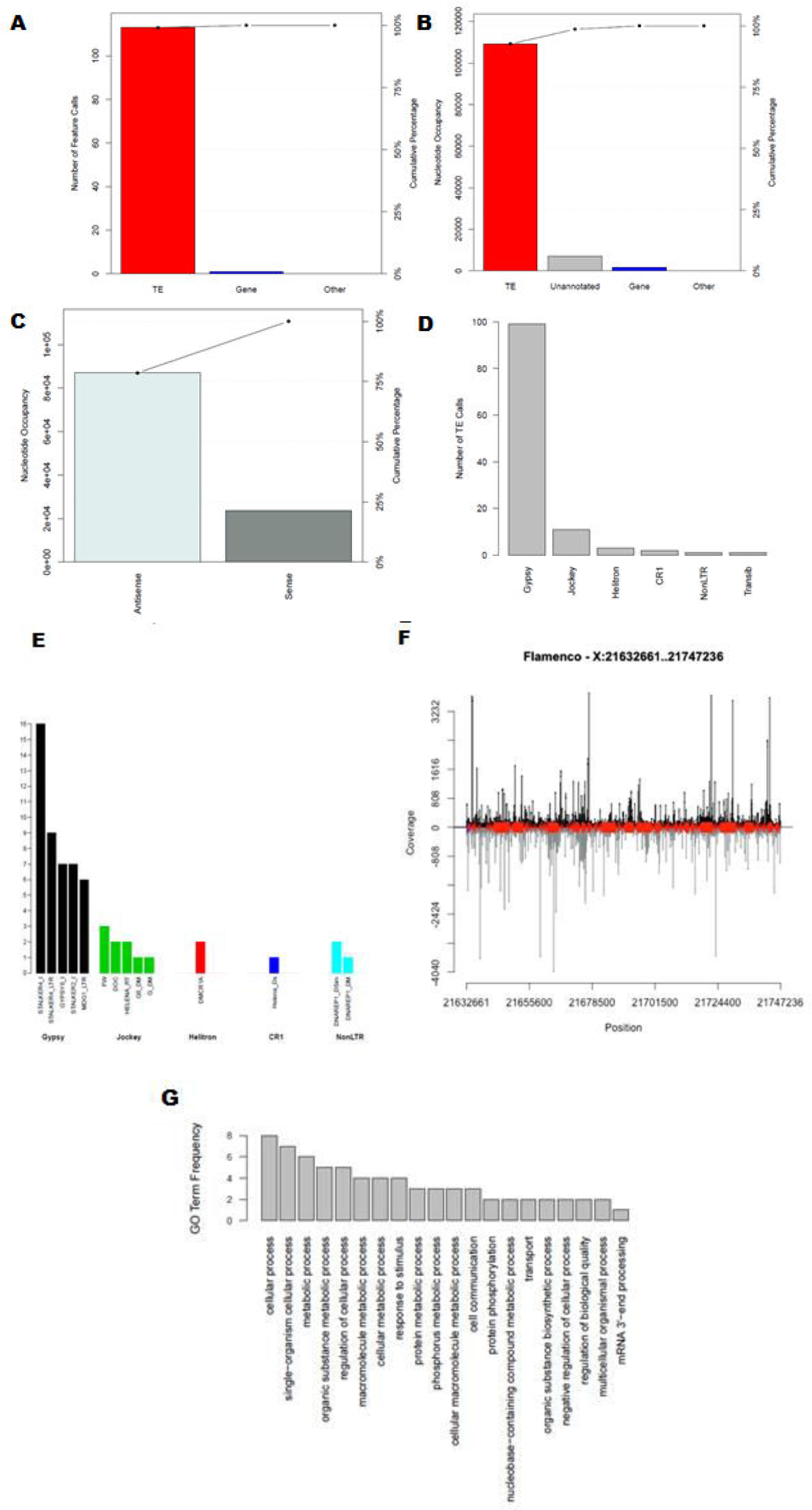
piRNA Cluster-Level Analysis of the *Flamenco* Locus of *Drosophila melanogaster*. **(A)** Number of known TE, Gene, or “Other” feature calls **(B)** Nucleotide occupancy of the feature calls **(C)** Orientation of feature calls **(D)** Number TE calls within the piRNA cluster for the most represented TE superfamilies **(E)** Number of individual TEs called in the top 5 represented superfamilies. Additional functionality can optionally be specified by the user to prompt production of a **(F)** sRNA coverage plot with feature content and orientation in 0-2hr eggs libraries **(G)** GO term frequency plot regarding all gene hits within the piRNA cluster.

## Results

### Benchmarking Software Performance

piClusterBusteR was timed for the analysis of the top 5 piRNA clusters identified in *Drosophila melanogaster* ovarian samples. When running sequentially on a single Intel(R) Xeon(R) CPU E5-2683 v4 at 2.10GHz, piClusterBusteR took approximately 3 hours to complete.

When utilizing the multithreading and multitasking capability of piClusterBusteR to analyze the same 5 piRNA clusters, using 5 nodes and 6 cores per node using the same processor speed, the timing of the piClusterBusteR run took approximately 20 minutes to run. One compute node was designated per piRNA cluster and six threads were utilized on each node. This run represents the enhanced capability of piClusterBusteR if additional resources, such as a computing cluster and queue submission system, are available to the user. Output from these independent analyses was identical.

piClusterBusteR results were observed on the previous established contents of the *flamenco* locus in *Drosophila melanogaster* which were extracted from the FlyBase database (Brennecke et al. 2007; Attrill et al. 2016).

Previous exploration with regard to the contents of the *flamenco* locus used RepeatMasker to characterize sequence content (Malone et al. 2009). The results of this method were extracted from the UCSC Table Browser retrieval tool (Karolchik et al. 2004). The contents and strand specificity of feature calls within the *flamenco* locus identified by piClusterBusteR were consistent with the previous observation (Figure 3.5).

**Figure 3.5.**
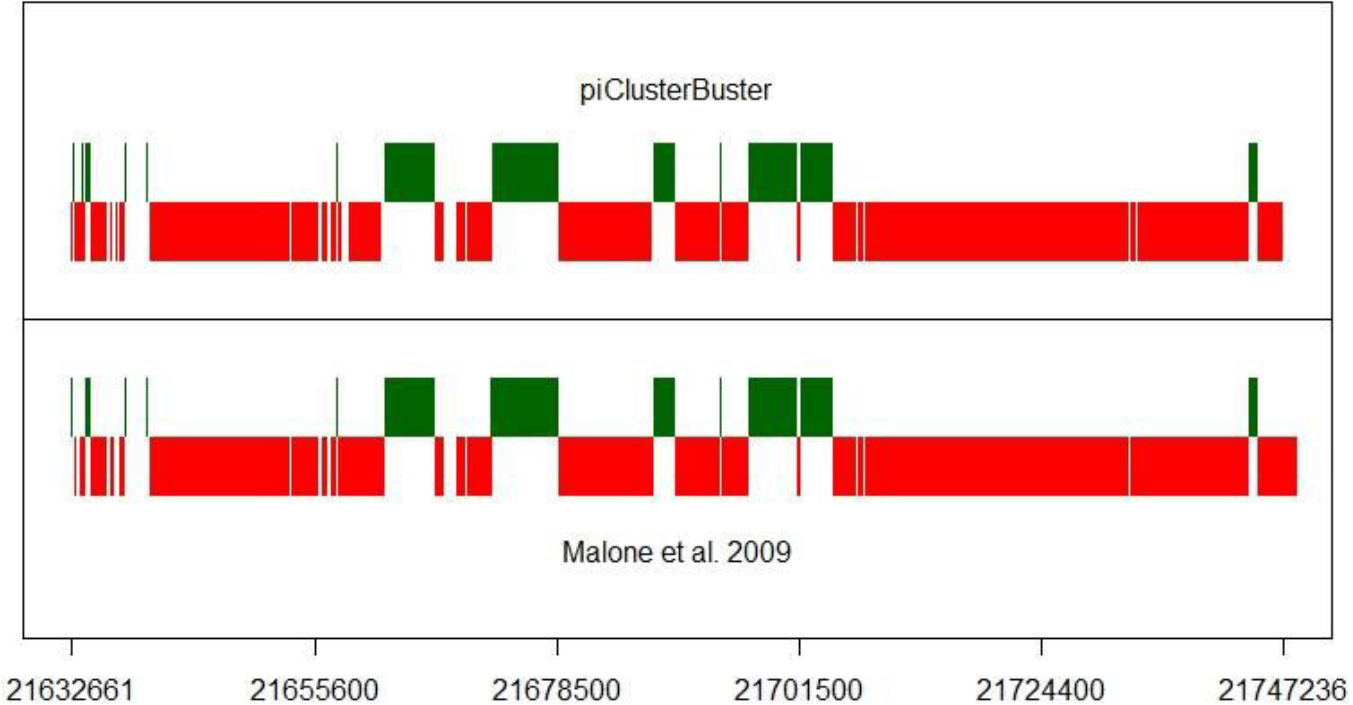
Comparison of Flamenco TE Annotation. A depiction of the agreement of piClusterBusteR characterization of the *flamenco* locus in comparison to previous reported characterization of TE contents in this locus in *Drosophila melanogaster* (Malone et al. 2009) (Figure 3.1S). Green boxes represent a sense orientation of TE calls and the red boxes represent an antisense orientation.

### piRNA Cluster Definition is Unaffected by Genome Size and Read Coverage

The 13 Metazoan species analyzed were selected based on data availability. A density-based approach of piRNA definition was implemented via use of the previous established software, proTRAC (Rosenkranz & Zischler 2012). A Pearson correlation test demonstrated that piRNA cluster definition by proTRAC appears to be irrespective of genome size of the organism and the number of piRNAs available for analysis at the 1% confidence level (Figure 3.6A-D, Figure 3.5S).

**Figure 3.6.**
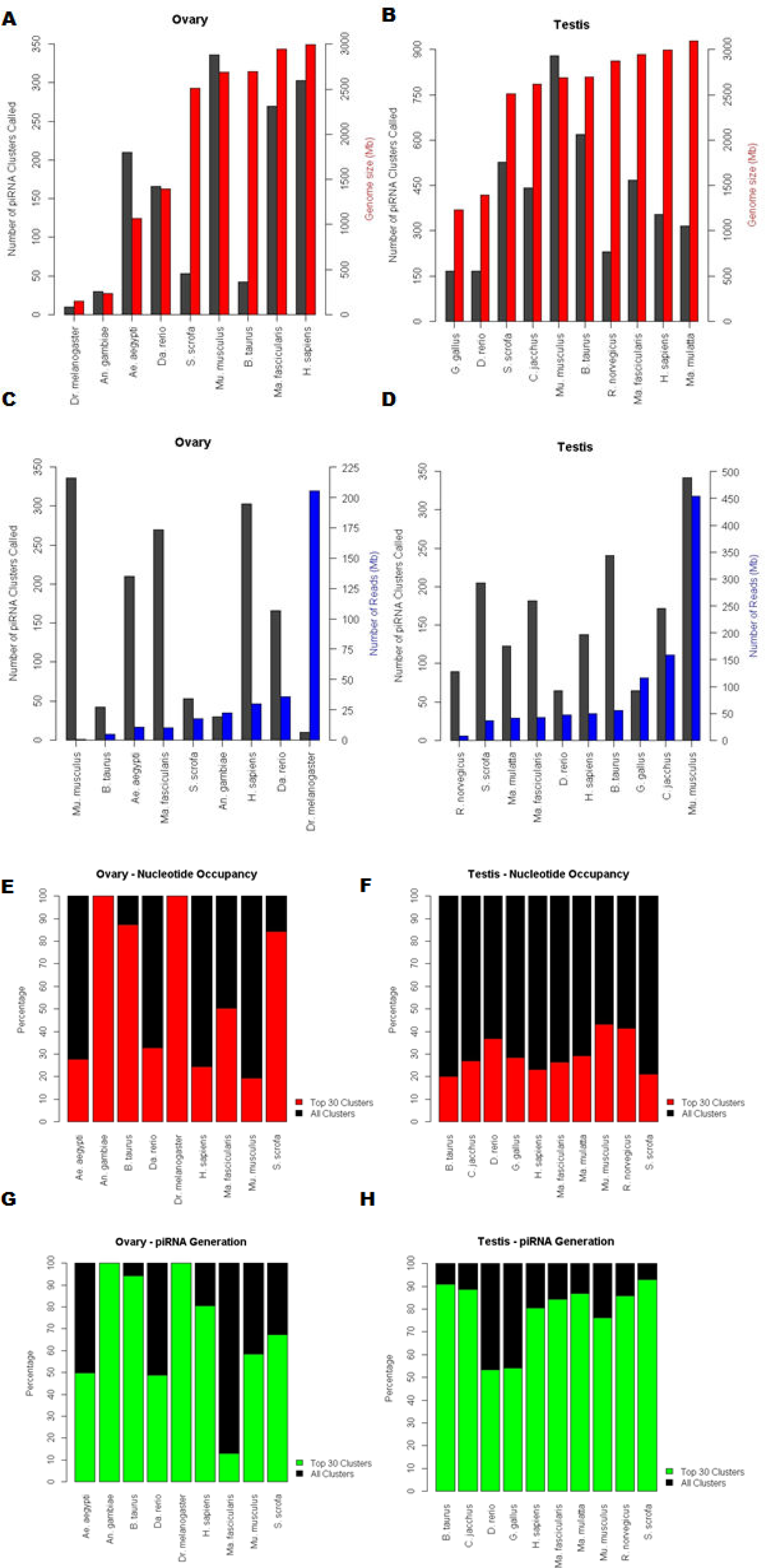
Representation of piRNA Cluster Loci. Number of piRNA cluster calls relative to the genome size of the organism in **(A)** ovary and **(B)** testis. Number of piRNA cluster calls relative to the number of reads available in **(C)** ovary and **(D)** testis. The percent composition of the top 30 piRNA clusters (red/blue) relative to the full repertoire of piRNA clusters (black) called by proTRAC with regard to the percent of nucleotides occupied by piRNA clusters in **(E)** ovary and **(F)** testis. The percentage of piRNA generated from the top piRNA cluster loci in **(G)** ovary and **(H)** testis

Given that the breadth of piRNA clusters is difficult to define in a given organism, due to the concern of false positives, we have used only the top 30 major contributing piRNA cluster loci in the between species comparisons of piRNA cluster composition. piRNA cluster definition required a length of at least five kilobases, at least 75% of the piRNAs deriving from a putative

piRNA cluster with a U-1 or A-10, at least 50% of the piRNAs deriving from a putative piRNA cluster with a U-1 and A-10, and the top 1% of piRNA sequences cannot comprise more than 90% of the piRNAs that were used to define a particular piRNA cluster.

In ovarian samples, the nucleotide occupancy of the top 30 piRNA clusters ranged from 19.3% to all of the piRNA clusters defined in a tissue with an average of 58.5% and median of 50.2% in these species (Figure 3.6E). The percent piRNA generation of the top 30 piRNA cluster loci ranged from 13.0% to all of the piRNAs generated in a tissue with an average of 68.0% and median of 67.3% relative to total piRNA generation (Figure 3.6G).

In testes samples, the nucleotide occupancy ranged from 20.1% to 43.1% relative to all of the piRNA clusters defined with an average of 29.7% and median of 27.8% in these species (Figure 3.6F). The percent piRNA generation of the top 30 piRNA cluster loci ranged from 53.3% to 93% with an average of 79.3% and median of 85.0% relative to total piRNA generation (Figure 3.6H).

Therefore, we consider the top 30 piRNA clusters to be representative of large scale architecture of genomic piRNA clusters based on the large proportion of the nucleotide occupancy and piRNA generation that is correlated with these loci.

### Top piRNA Cluster Architecture is Conserved in Metazoans on a Large Scale

The analysis of piRNA cluster architecture focused on the number of piRNA clusters, piRNA cluster size, the known features within the piRNA cluster, and the orientation of the known feature.

Certain features of piRNA cluster architecture were conserved better than others. In all of the Metazoan species observed in this analysis, the majority of piRNA cluster sequence was unable to be attributed to any known origin. Unannotated sequence ranged between 18% and 70% of piRNA cluster composition. TEs were the major known contributor to piRNA cluster loci. TEs occupied up to 78% of ovarian piRNA cluster loci and 62% of testis piRNA clusters, with an average piRNA cluster occupancy of 40% to 32% in ovarian and testes libraries, respectively. Sequences of known genic origin ranged from 1 to 11%, with an average of 3% and 3.5% piRNA cluster occupancy in ovaries and testes, respectively. Non-genic, non-TE sequences within the NCBI database were the least significant contributor to piRNA cluster loci in these species, ranging from 0 to 9% of piRNA cluster composition, with an average piRNA cluster occupancy between 4 to 3.5% in ovarian and testes libraries (Figure 3.7).

**Figure 3.7.**
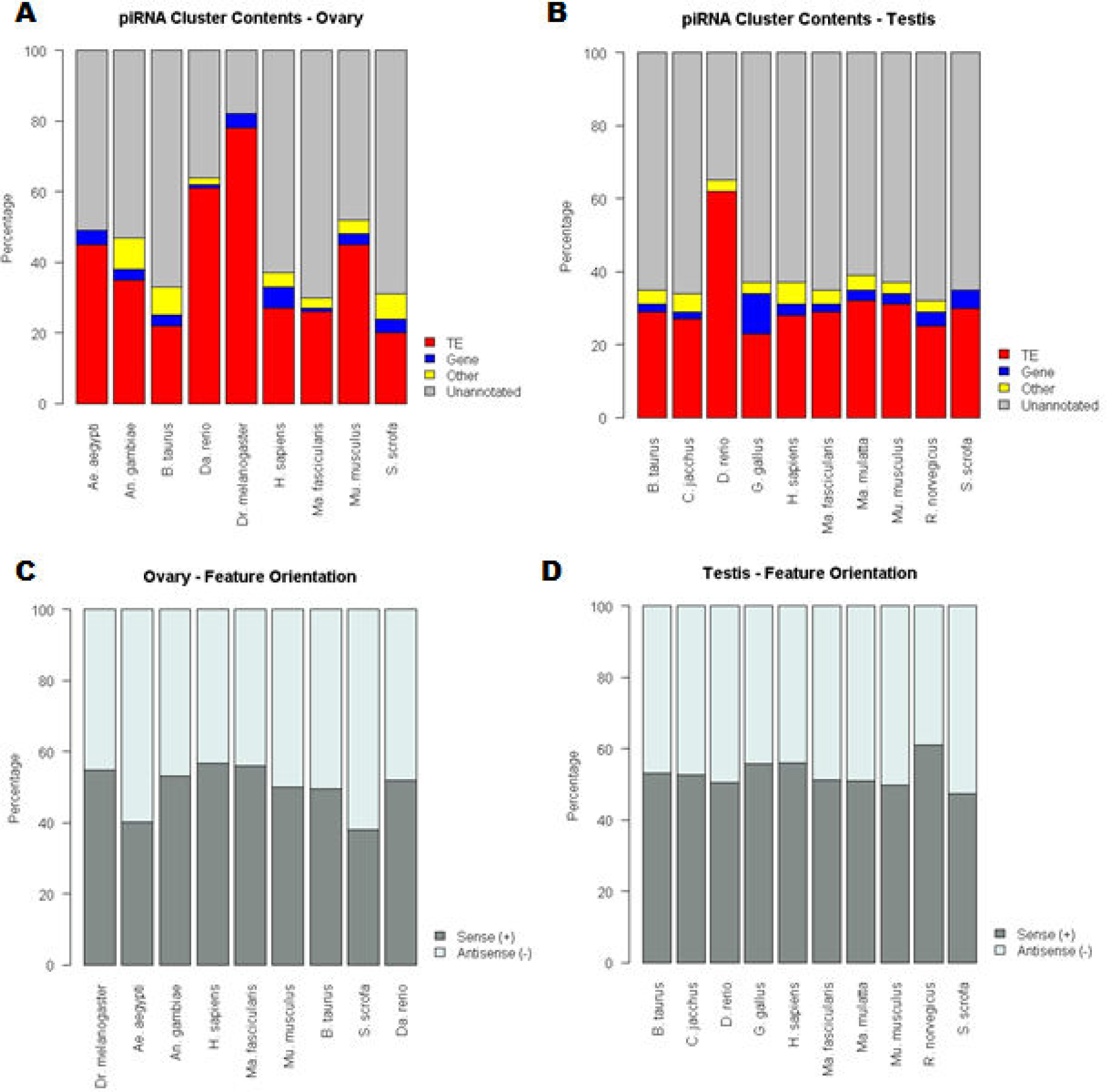
Comparison of Top piRNA Cluster Composition. piRNA cluster content comparison between species in **(A)** ovary and **(B)** testis. Orientation of the feature calls in **(C)** ovary and **(D)** testis samples. These data represent an analysis of the top 30 piRNA cluster loci in each species. Available ovarian and testes datasets from the piRNA cluster database and the Short Read Archive were used to run piClusterBusteR (Rosenkranz 2016; Kodama et al. 2012). Only ten piRNA clusters were called in the *Dr. melanogaster* covarian library

The strand specificity of feature calls within top piRNA cluster loci was also summarized. Features were predominantly characterized on the sense strand of piRNA clusters. The nucleotide occupancy of sense features accounted for between 38.0% to 61.0% of feature calls with a piRNA cluster with an average of 50.0% and 52.8% in ovarian and testes samples, respectively, in these species.

### Tissue-Specificity of piRNA Cluster Loci

piRNA cluster definition can vary between sRNA libraries that derived from the same tissue. The number of defined piRNA clusters differed from 3 to 398 calls between two samples of the same tissue with a mean difference of 75 and median difference of 45 piRNA cluster calls. Although, at least 52.4%, and up to 92.2% of the lesser piRNA cluster definitions were also represented in the larger sample of piRNA cluster calls. piRNA cluster definition demonstrated an average of 67.8% overlap in same tissue samples (Figure 3.8).

**Figure 3.8.**
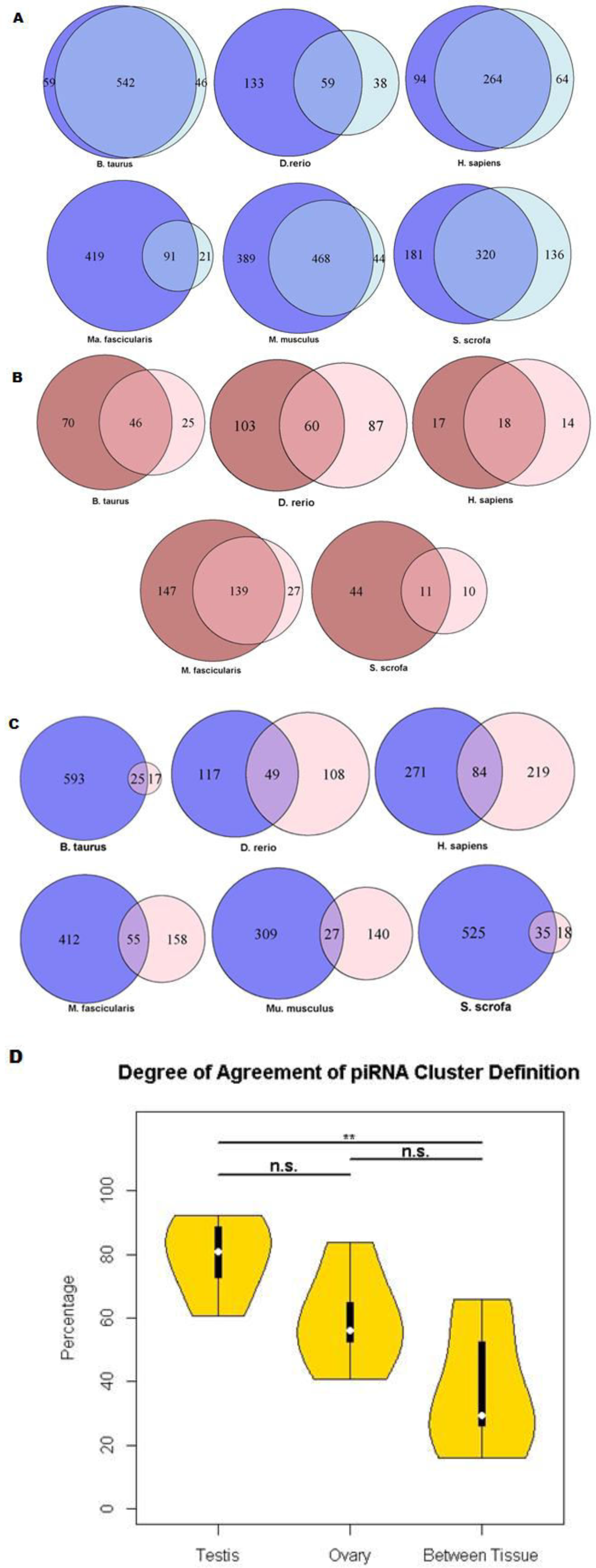
Tissue-specificity of piRNA Cluster Definition. Venn Diagrams representing the degree of overlap of piRNA cluster definitions between two independent libraries of the same tissue. **(A)** Blue circles represent testes samples and **(B)** red circles represent ovarian samples. Only one ovarian sRNA library was analyzed in *M. musculus* and *D. melanogaster* testis defined no piRNA clusters. **(C)** Venn Diagrams representing the degree of overlap of piRNA cluster definitions between ovary and testis piRNAs. **(D)** Violin plot representing the agreement of piRNA cluster calls within same species testes, within same species ovaries, and a comparison between testes and ovaries piRNA cluster definition. (Figure 3.12S)

We observed a significantly lesser degree of agreement piRNA cluster calls relative to same sample testis libraries. Same sample ovary libraries also showed a small increase in piRNA cluster agreement relative to samples between tissues (Figure 3.8D). The number of piRNA cluster definition differed from 9 to 576 calls between two samples of the same tissue with a mean difference of 255 and median difference of 211 piRNA cluster calls. The lesser sample of piRNA cluster definitions ranged in agreement from 16.2% to 66.0% with an average of 34.6% agreement and a median of 29.5% between samples (Figure 3.8C).

## Discussion

The options for piClusterBusteR performance enhancement allow for utilization of multitasking and multithreading. Use of multithreading allows for the execution of piClusterBusteR processes by multiple nodes simultaneously. Utilization of the multitasking capability of piClusterBusteR prompts independent, parallel submission for each piRNA cluster of interest to independent compute nodes. Multithreading and multitasking piClusterBusteR runs allows the user the capability to significantly increase the number of piRNA clusters under observation without significantly increasing the timing of the piClusterBusteR run. piClusterBusteR supports both Torque/Maui or Slurm resource management software.

This comparison of piRNA cluster architecture focuses on the major genomic loci contributing to piRNA populations. By only considering only the top piRNA cluster loci in this analysis, we can be relatively confident in piRNA cluster definition relative to other piRNA-generating loci. The top 30 piRNA clusters also were a large representation of the total nucleotides occupied by piRNA clusters in these genomes, as well as disproportionally large contributors to total piRNA populations in these species (Figure 3.6). Taken together, the contents of the piRNA clusters in the analysis serve as the best representation of piRNA cluster architecture in these species.

Since it is difficult to determine whether RepeatMasker and CENSOR will annotate a piRNA cluster of interest more thoroughly, with higher confidence, we implemented nested annotation. A nested annotation approach allows for both of the programs that performed well in annotating sequences that are dense with repeats, RepeatMasker and CENSOR, the opportunity to characterize the sequence of interest, while only maintaining the best annotation in the description of the contents of piRNA cluster sequence (Figure 3.2) (Smit et al. 1996; Jurka et al. 1996). This method allows for consistent and accurate characterization amongst diverse piRNA clusters on a large scale.

We also noted that the degree of sense or antisense orientation of feature calls within individual piRNA clusters correlated with the direction of transcription in known piRNA clusters, *flamenco* and 42AB (Brennecke et al. 2007; Malone et al. 2009). Therefore, the orientation of feature calls within a piRNA cluster may be informative in the prediction of the nature of piRNA cluster transcription.

Components of piRNA architecture were strikingly similar across species. With regard to known piRNA cluster features, TEs consistently composed the majority by nucleotide occupancy and a relatively low percentage of known genic and “other” calls. The majority of informative, “other” hits within the NCBI nucleotide database were associated with mRNAs that were not available in the organism-specific gene set. Other informative non-genic, non-TE sequence appeared to be of viral and rRNA origin. The orientation of feature calls within piRNA clusters were also highly conserved in these species. Taken together, these data suggest highly conserved nature, yet dynamic capacity within piRNA cluster architecture with regard to known features in Metazoans (Figure 3.6).

We observed that a significant portion of the piRNA cluster sequence was unable to be characterized in the species observed in this study (Figure 3.7). This observation prompts an interesting question regarding the derivation of piRNA cluster sequence whose origin is currently undetectable and its purpose within the piRNA clusters. This sequence is of particular biological interest given that these sequences occupy significant regions of piRNA clusters and may further inform scientific knowledge of piRNA cluster biogenesis and function.

Sets of piRNA clusters were differentially represented between different, and within the same, independent tissue samples. It is worth noting that differential representation of piRNA generating loci between same, independent tissue samples may be due to the previous observation that sRNA libraries represent only a subset of the complete sRNA populations within the cell, even when deep sequencing is performed (Yamtich et al. 2015). However, the variability in piRNA cluster overlap was far greater between tissues than within tissues when comparing piRNA cluster definitions between libraries. Therefore, preliminary observation of these data supports a model in which different regions of the genome appear to be responsible for generating the majority of piRNAs in ovaries and testes samples in Metazoans and it may be advantageous for an organism to have a diverse, dynamic set of piRNA cluster activity in a unique cellular environment.

